# Albendazole reduces endoplasmic reticulum stress induced by *Echinococcus multilocularis* in mice

**DOI:** 10.1101/2021.02.03.429530

**Authors:** Michael Weingartner, Fadi Jebbawi, Junhua Wang, Simon Stücheli, Bruno Gottstein, Guido Beldi, Britta Lundström-Stadelmann, Alex Odermatt

**Author notes:** To whom correspondence should be addressed: Dr. Alex Odermatt, Division of Molecular and Systems Toxicology, Department of Pharmaceutical Sciences, University of Basel, Klingelbergstrasse 50, 4056 Basel, Switzerland. These authors contributed equally to the presented study (Michael Weingartner, Fadi Jebbawi). **Disclosures:** The authors declare no commercial or financial conflict of interest.

## Abstract

**Background:** *Echinococcus multilocularis* causes alveolar echinococcosis (AE), a rising zoonotic disease in the northern hemisphere. Treatment of this fatal disease is limited to chemotherapy using benzimidazoles and surgical intervention, with relatively frequent disease recurrence in cases without radical surgery. Elucidating the molecular mechanisms underlying *E. multilocularis* infections and host-parasite interactions aids developing novel therapeutic options. This study explored an involvement of unfolded protein response (UPR) and endoplasmic reticulum-stress (ERS) during *E. multilocularis* infection in mice.

**Methods:** *E. multilocularis-* and mock-infected C57BL/6 mice were subdivided six weeks after infection into vehicle and albendazole (ABZ) treated groups. Eight weeks later, liver tissue was collected to examine mRNA, microRNA (miR) and protein expression of UPR- and ERS-related genes.

**Results:** *E. multilocularis* infection upregulated UPR- and ERS-related proteins, including ATF6, CHOP, GRP78, ERP72, H6PD and calreticulin, whilst PERK and its target eIF2α were not affected, and IRE1α and ATF4 were downregulated. ABZ treatment in *E. multilocularis* infected mice reversed the increased ATF6 and calreticulin protein expression, tended to reverse increased CHOP, GRP78, ERP72 and H6PD expression, and decreased ATF4 and IRE1α expression to levels seen in mock-infected mice. The expression of miR-146a-5p (downregulated by IRE1α) and miR-1839-5p (exhibiting a unique target site in the IRE1α 3’UTR) were significantly increased in *E. multilocularis* infected mice, an effect reversed by ABZ treatment. Other miRs analyzed were not altered in *E. multilocularis* infected mice.

**Conclusions and Significance:** AE causes UPR activation and ERS in mice. The *E. multilocularis*-induced ERS was ameliorated by ABZ treatment, indicating its effectiveness to inhibit parasite proliferation and downregulate its activity status. ABZ itself did not affect UPR in control mice. Identified miR-146a-5p and miR-1839-5p might represent biomarkers of *E. multilocularis* infection. Modulation of UPR and ERS, in addition to ABZ administration, could be exploited to treat *E. multilocularis* infection.

**Author summary:** Alveolar echinococcosis is a zoonotic disease caused by the fox tapeworm *Echinococcus multilocularis*. Treatment of this fatal disease is limited to surgical intervention, preferably radical curative surgery if possible, and the use of parasitostatic benzimidazoles. It is not yet fully understood how the parasite can remain in the host’s tissue for prolonged periods, complicating the development of therapeutic applications. This work investigated an involvement of the unfolded protein response (UPR) and endoplasmic reticulum-stress (ERS) during *E. multilocularis* infection and upon treatment with albendazole (ABZ) in mice. The results revealed increased expression levels of the ERS sensor ATF6 and of downstream target genes in liver tissue of *E. multilocularis-* compared to mock-infected mice. Additionally, H6PD, generating NADPH within the endoplasmic reticulum, and the lectin-chaperone calreticulin were increased in *E. multilocularis* infected liver tissue while the expression of the ERS associated genes ATF4 and IRE1α were decreased. The miR-1839-5p and miR-146-p, linked to IRE1α, were elevated upon *E. multilocularis* infection, offering potential as novel biomarkers of alveolar echinococcosis. The observed gene expression changes were at least partially reversed by ABZ treatment. Whether modulation of UPR and ERS targets can improve the therapy of alveolar echinococcosis remains to be investigated.

## Introduction

Alveolar echinococcosis (AE) is a severe helminthic disease caused by accidental ingestion of eggs from the fox tapeworm *Echinococcus multilocularis (E. multilocularis)* [1, 2]. After an incubation period of 5 up to 15 years without perceivable symptoms, AE has a fatal outcome in up to 90% of cases when left untreated [3-5]. AE is characterized by a slow but progressive tumor-like growth of metacestodes (larval stage) mainly in the liver, with a tendency to spread to various organs like spleen, brain, heart and other tissues such as bile ducts and blood vessels [6-8]. Treatment by radical surgical resection is limited by the diffuse infiltrations of AE lesions in liver and other tissues in advanced cases [9, 10]. If lesions cannot be completely removed by surgery, a lifelong medication is required, usually using benzimidazoles, which can cause adverse side effects. For example, several cases with hepatotoxic effects due to treatment with the benzimidazole albendazole (ABZ) were reported with various outcomes [11-14]. An inadequate adherence to chemotherapy, due to adverse side effects, can explain the relapsing spread of AE and a worsening general condition of patients with severe *E. multilocularis* infiltrations [15, 16]. Considering these circumstances, the rising numbers of reported cases of AE especially in Europe and the lack of a curative drug treatment, emphasizes the necessity to further investigate the mechanisms underlying this threat and search for improved therapeutic options [17-22].

Several bacteria and viruses have been described to modulate endoplasmic reticulum (ER) stress (ERS) and unfolded protein response (UPR), either by bacterial virulence factors such as toxins (e.g. cholera toxin, pore-forming toxins) or by the increased demand of newly synthesized proteins for the production of virions [23-28]. Activation of the UPR via an induction of glucose-regulated protein 78 (GRP78) has previously been shown in cells infected with *Human immunodeficiency virus* (HIV) [27, 29], *Dengue virus* (DENV) [30], *West Nile virus* (WNV) [31] or *Human cytomegalovirus* (HCMV) [32]. Moreover, facilitated replication of viruses and immune evasion represent key features following UPR activation by *Mouse hepatitis virus* (MHV) [33] and *Herpes simplex virus 1* (HSV-1) [34]. On the other hand, an ERS-induced upregulation of UPR-related genes was linked to an enhanced production of pro-inflammatory cytokines in B-cells and stellate cells [35, 36]. A modulation of the UPR pathway was reported not only during viral but also bacterial infections. *Legionella pneumophila* infection led to an inhibition of X-box binding protein 1 (XBP1) splicing in mammalian host cells, thereby suppressing the host UPR pathway [37]. *Mycobacterium tuberculosis* was found to induce ERS, indicated by increased CCAAT/enhancer-binding protein homologous protein (CHOP) and GRP78 protein levels in infected macrophages, leading to host cell apoptosis. Decreased levels of phosphorylated eukaryotic initiation factor 2α (eIF2α) in infected cells were associated with enhanced bacterial survival [38].

However, to date the knowledge of pathogen-induced ERS and UPR activation is incomplete and mainly limited to bacterial and viral infections. In this regard, a modulation of the host’s UPR with an upregulation of CHOP was observed in *Toxoplasma gondii* infected cells, leading to apoptosis of host cells [39]. Another study in a mouse model provided evidence that *Plasmodium berghei* exploits the host’s UPR machinery for its survival [40]. However, little is known about the involvement of ERS and UPR activation in *E. multilocularis* infection.

The present study addressed a possible role of the modulation of UPR- and ERS-related proteins in host cells in response to AE and investigated the effect of these pathways using *E. multilocularis* infected mice as a model. A better understanding of a contribution of proteins of the UPR and ERS pathways in the context of infectious diseases is of interest regarding the development of improved therapeutic strategies to cope with parasitic infections [41-43].

Additionally, this work investigated whether microRNAs (miRs), small non-coding single stranded RNAs (17-24 nucleotides) that regulate the post-transcriptional levels of mRNAs by inhibiting their translation to proteins, may be altered upon *E. multilocularis* infection. Several studies revealed a functional interaction between UPR/ERS signaling and the expression of miRs [44-46]. Silencing of miRs was found to be involved in ERS signaling and miRs act as effectors and modulators of the UPR and ERS pathways [47]. The miRs, isolated from human specimen, including urine, saliva, serum and tissues, are considered as biomarkers of several immune pathologies such as cancer, autoimmune diseases and viral or bacterial infections [48-54]. In this regard, recent investigations provided evidence for a role of some miRs in the regulation of UPR signaling, with miR-181a-5p and miR-199a-5p shown to suppress the UPR master regulator GRP78 [47, 55, 56]. On the other side, UPR pathways also can affect the expression of some miRs, as shown by inositol-requiring enzyme 1α (IRE1α) that cleaves the precursors of anti-apoptotic miR-17-5p, miR-34a-5p, miR-96-5p and mir-125b-5p, which in turn negatively regulate the expression of caspase 2 and thioredoxin-interacting protein [57, 58]. In addition, the activation of protein kinase R (PKR)-like ER kinase (PERK) induces the expression of miR-30c-2-3p, which downregulates XBP1, representing a possible negative crosstalk between PERK and IRE1α [58].

Boubaker et al. recently described a murine miR signature in response to early stage *E. multilocularis* egg infection where the expression of seven miRs (miR-148a-3p, miR-143-3p, miR-101b-3p, miR-340-5p, miR-22-3p, miR-152-3p and miR-30a-5p) was decreased in AE-infected compared to mock-infected mice. In contrast, *E. multilocularis* infected mice exhibited significantly higher levels of miR-21a-5p, miR-28a-5p, miR-122-5p and miR-1839-5p compared to the mock-infected controls [59]. The miRs mentioned above were therefore also analyzed in the present study.

## Materials and Methods

### Chemicals and reagents

Polyvinylidene difluoride (PVDF) membranes (Cat# IPVH00010, pore size: 0.45 µm), Immobilon Western Chemiluminescence horseradish-peroxidase (HRP) substrate kit, radioimmunoprecipitation assay (RIPA) buffer, β-mercaptoethanol, HRP-conjugated goat anti-mouse secondary antibody (Cat# A0168, RRID:AB_257867), rabbit polyclonal anti-hexose-6-phosphate dehydrogenase (H6PD) antibody (Cat# HPA004824, RRID:AB_1079037), protease inhibitor cocktail, dNTPs, KAPA SYBR® FAST kit and qPCR kit were purchased from Merck (Darmstadt, Germany). RNeasy Mini kit and QIAcube were obtained from Qiagen (Venlo, Netherlands), GoScript reverse transcriptase (Cat# A5003) from Promega (Fitchburg, WI, USA), rabbit monoclonal anti-lamin B1 antibody (Cat# ab133741, RRID:AB_2616597) from Abcam (Cabridge, UK) and mouse monoclonal anti-GRP78 antibody (Cat# 610978, RRID:AB_398291) from BD Bioscience (Franklin Lakes, NJ, USA). HRP-conjugated goat anti-rabbit secondary antibody (Cat# 7074, RRID:AB_2099233), mouse monoclonal anti-CHOP antibody (Cat# 2895, RRID:AB_2089254), rabbit polyclonal anti-calreticulin (CRT) antibody (Cat# 2891, RRID:AB_2275208), rabbit polyclonal anti-eIF2α antibody (Cat# 9722, RRID:AB_2230924), rabbit monoclonal anti-ATF4 antibody (Cat# 11815, RRID:AB_2616025) and rabbit monoclonal anti-ATF6 antibody (Cat# 65880, RRID:AB_2799696) were purchased from Cell Signaling (Cambridge, UK). Mouse monoclonal anti-PERK antibody (Cat# sc-377400, RRID:AB_2762850), anti-IRE1α antibody (Cat# sc-390960, RRID: N/A) and anti-ERp72 antibody (Cat# sc-390530, RRID: N/A) were obtained from Santa Cruz Biotechnology (Dallas, TX, USA). Pierce® bicinchoninic acid protein assay kit, Nanodrop™ One C (Cat# 13-400-519) and Trizol® total RNA isolation reagent were purchased from Thermo Fisher Scientific (Waltham, MA, USA). Precellys-24 tissue homogenizer was purchased from Bertin Instruments (Montigny-le-Bretonneux, France). Primers for real-time quantitative polymerase chain reaction (RT-qPCR) were obtained from Microsynth (Balgach, Switzerland). TaqMan microRNA Assays, snoRNA234, TaqMan microRNA reverse transcription kit (Cat# 4366596), TaqMan fast advanced master mix (Cat# 4444556), TaqMan probes (Cat# 4427975, Assay IDs 000468, 000389, 000398, 000470, 121135_mat, 000416, and 001234) and ViiA 7 real-time PCR system (Cat# 4453545) were purchased from Applied Biosystems (Foster City, CA, USA). Rabbit polyclonal anti-calnexin (CNX) antibody (Cat# SAB4503258, RRID:AB_10746486) and all other reagents were purchased from Sigma-Aldrich (St. Louis, MO, USA).

### Ethics Statement

The animal studies were performed in compliance with the recommendations of the Swiss Guidelines for the Care and Use of Laboratory Animals. The protocol used for this work was approved by the governmental Commission for Animal Experimentation of the Canton of Bern (approval no. BE112/17).

### Animal experimentation and sampling

Animal experimentation, liver tissue extraction and corresponding liver tissue samples were previously described by Wang et al. [60]. Briefly, female 8-week-old wild type C57BL/6 mice were randomly distributed into 4 groups with 6 animals per group: 1) mock-infected (corn oil treated) control mice (referred to as “CTRL”); 2) *E. multilocularis* infected, vehicle treated mice (referred to as “AE”); 3) *E. multilocularis* infected, ABZ-treated mice (referred to as “AE-ABZ”); and 4) mock-infected, ABZ-treated mice (referred to as “ABZ”) (S1 Fig). All animals were housed under standard conditions in a conventional daylight/night cycle room with access to feed and water *ad libitum* and in accordance with the Federation of European Laboratory Animal Science Association (FELASA) guidelines. During the experimental period animals were examined weekly for subjective presence of health status and changes in weight. At the end of the experiment the mice were euthanized by CO_2_ and liver tissue was resected followed by immediate freezing in liquid nitrogen and storage at −80°C until use.

### Parasite preparation and secondary infection of mice by intraperitoneal administration

Infection with *E. multilocularis* by intraperitoneal injection was conducted as previously described [61]. Briefly, *E. multilocularis* (isolate H95) was extracted and maintained by serial passages in C57BL/6-mice. Aseptic removal of infectious material from the abdominal cavity of infected animals was used for continuation of AE in mice. Collected tissue was grinded through a sterile 50 μm sieve, roughly 100 vesicular cysts were suspended in 100 μL sterile PBS and administrated *via* intraperitoneal injection to group 2 (“AE”) and 3 (“AE-ABZ”). Mice of the mock-infected groups 1 (“CTRL”) and 4 (“ABZ”) received 100 μL of sterile PBS.

### Treatment

Treatment started 6 weeks after initial infection (S1 Fig). 100 μL corn oil were orally administrated to groups 1 (“CTRL”) and 2 (“AE”) five times per week. Group 3 (“AE-ABZ”) and 4 (“ABZ”) received 100 μL corn oil containing ABZ (200 mg/kg body weight) orally five times per week. The treatment was terminated after 8 weeks and mice were euthanized.

### Analysis of protein expression by western blot

The procedures for liver sample preparation and western blot analysis have been previously described [62]. Briefly, liver samples (approximately 7 mg) were homogenized (30s, 6500 rpm, at 4°C, using a Precellys-24 tissue homogenizer) in 450 µL RIPA buffer (50 mM Tris-HCl, pH 8.0, with 150 mM NaCl, 1.0% NP-40, 0.5% sodium deoxycholate and 0.1% sodium dodecyl sulfate) containing protease inhibitor cocktail and centrifuged (4 min, 16,000 × g, 4°C). Protein concentration was measured by a standard bicinchoninic acid assay (Pierce® BCA Protein Assay Kit). Samples were boiled (5 min at 95°C) in Laemmli solubilization buffer (60 mM Tris-HCl, 10% glycerol, 0.01% bromophenol blue, 2% sodium dodecyl sulfate, pH 6.8, 5% β-mercaptoethanol). The protein extract (20 μg) was separated by 10-14% SDS-PAGE and blotted on PVDF membranes. The membranes were blocked (1 h, room temperature) in TBST-BSA, (20 mM Tris buffered saline with 0.1% Tween-20, 1% bovine serum albumin). All primary and secondary antibody dilutions and incubations were performed in TBST-BSA. For the detection of primary antibodies raised in rabbit, secondary HRP-conjugated goat anti-rabbit antibody was used. Primary antibodies raised in mouse were detected by HRP-conjugated goat anti-mouse antibody. Primary antibodies were incubated at 4°C over-night. Secondary antibodies were applied at room temperature for 1 h. Protein content was visualized by Immobilon Western Chemiluminescence HRP substrate. Protein bands were quantified by densitometry normalized to Lamin B1 protein levels using ImageJ software (version 1.53n). The applications of primary and secondary antibodies can be found in S1 Table.

### Quantification of mRNA by RT-qPCR

Methods for preparation of liver samples, RNA isolation and RT-qPCR analysis were performed as described [60]. Briefly, total RNA was isolated from liver tissue (approximately 8 mg) by homogenization (30 s, 6500 rpm, 4°C; Precellys-24 tissue homogenizer) in 350 µL RLT buffer (RNeasy Mini Kit) supplied with 40 mM dithiothreitol, followed by centrifugation (3 min, 16,000 × g). The supernatant was further processed according to the manufacturer’s protocol for RNA isolation from animal tissues and cells using QIAcube. RNA quality and concentration was analyzed using Nanodrop™ One C. 1000 ng of RNA was transcribed into cDNA using GoScript Reverse Transcriptase. KAPA SYBR^®^ FAST Kit was used for RT-qPCR (4 ng of cDNA per reaction in triplicates, 40 cycles) analysis, and reactions were performed using a Rotor Gene Real-Time Cycler (Corbett Research, Sydney, New South Wales, Australia). Data was normalized to the expression levels of the endogenous control gene β-actin. Comparison of gene expression was performed using the 2-ΔCT-method using β-actin as housekeeping gene [63]. Primers used for RT-qPCR are listed in S2 Table.

### Extraction and quantification of miRNA by qPCR

Total RNA was extracted from liver tissues using Trizol® total RNA isolation reagent and RNA concentration quantified using Nanodrop™ One C. TaqMan microRNA assays were used to quantify mature miR expression. SnoRNA234 was used as endogenous control of miR expression. Thus, miR-specific reverse transcription was performed for each miR using 10 ng of purified total RNA, 100 mM dNTPs, 50 U multiple reverse transcriptase, 20 U RNase inhibitor, and 50 nM of miR-specific reverse transcription primer samples using the TaqMan MicroRNA Reverse Transcription kit. Reactions with a volume of 15 µL were incubated for 30 min at 16°C, 30 min at 42°C, and 5 min at 85°C to inactivate the reverse transcriptase. RT-qPCR (5 µL of reverse transcription product, 10 µL TaqMan Fast Advanced Master Mix and 1 µL TaqMan microRNA Assay Mix containing PCR primers and TaqMan probes) were run in triplicates at 95°C for 10 min followed by 40 cycles at 95°C for 15 s and 60°C for 1 min. Quantitative miR expression data were acquired and analyzed using the ViiA 7 real-time PCR system (Applied Biosystems, Cat# 4453545).

### Statistical analyses

Data are presented as mean ± SD. The significance of the differences between the examined animals were determined by Kruskal-Wallis test or one-way ANOVA, whereby the specific test is indicated in the Figure legend. No outliers were excluded. *P≤0.05; **P≤0.01; ***P≤0.001 significantly different as indicated. GaphPad Prism software (version 8.0.2, GraphPad, La Jolla, CA, USA) was used for statistical analysis.

## Results

### Effects of AE on the expression of proteins related to UPR and ERS pathways

As the present knowledge on the modulation of UPR and ERS pathways during parasitic infections is limited, this study examined the expression of key proteins related to these pathways in liver tissues of mice infected with *E. multilocularis*. In the applied model of secondary *E. multilocularis* infection, differential effects on the expression of proteins of the different UPR and ERS branches were observed. Among the PERK pathway, ATF4 protein levels were significantly decreased in liver tissue of AE mice compared to mock-infected controls (Table 1, Fig 1). The expression of PERK and its target protein eIF2α were not affected by *E. multilocularis* infection. However, the most pronounced effects were observed for ERS related proteins of the ATF6 branch of the UPR (Table 1, Fig 2). The levels of all four proteins analyzed were elevated, whereby the luminal chaperone and protein disulfide isomerase ERP72 and the ERS marker CHOP were 2.0-fold and 4.5-fold increased and ATF6 and GRP78 tended to be elevated with 2.2-fold and 2.7-fold higher levels, respectively. IRE1α protein expression was decreased by about 3-fold in *E. multilocularis* infected compared to control mouse liver tissues (Table 1, Fig 3), whilst its target the spliced form of XBP1 showed a trend to lower levels in liver tissues of infected mice (S2 Fig). However, XBP1 protein expression could not be assessed as no specific antibody could be identified.

**Table 1.** Expression of proteins involved in UPR and ERS pathways. Protein levels in liver tissue samples were analyzed by western blot and densitometry (animals per group n=6). Numbers represent protein expression levels normalized to those of the control (CTRL) group (mean ± SD). Significantly decreased protein levels are highlighted in red and increased protein levels in blue. Symbols indicate significant differences (p≤0.05) between groups: *, compared to CTRL; §, compared to ABZ; #, compared to AE-ABZ. No outliers were excluded. Non-parametric, Kruskal-Wallis test.

|  |  | Group |  |  |  |
| --- | --- | --- | --- | --- | --- |
| Classification Pathway | Relative protein expression (normalized to CTRL) | CTRL (n=6) | AE (n=6) | AE-ABZ (n=6) | ABZ (n=6) |
| PERK branch | PERK | 1.0 | 1.1 ($\pm 0.8$) | 1.1 ($\pm 0.5$) | 2.8 ($\pm 2.6$) |
| | eIF2α | 1.0 | 1.5 ($\pm 0.5$) | 1.6 ($\pm 0.4$) | 1.1 ($\pm 0.4$) |
| | ATF4 | 1.0 | 0.3 ($\pm 0.1$)*,§ | 0.8 ($\pm 0.2$) | 1.0 ($\pm 0.4$) |
| ATF6 branch | ATF6 | 1.0 | 2.2 ($\pm 1.0$)# | 0.6 ($\pm 0.2$) | 1.2 ($\pm 0.5$) |
| | CHOP | 1.0 | 4.5 ($\pm 2.0$)*,§ | 3.0 ($\pm 1.4$)§ | 1.0 ($\pm 0.3$) |
| | GRP78 | 1.0 | 2.7 ($\pm 1.5$)§ | 1.3 ($\pm 1.1$) | 1.0 ($\pm 0.3$) |
| | ERP72 | 1.0 | 2.0 ($\pm 0.6$)*,§ | 1.3 ($\pm 0.6$) | 0.9 ($\pm 0.4$) |
| IRE1 branch | IRE1α | 1.0 | 0.3 ($\pm 0.2$)*,§ | 0.8 ($\pm 0.4$) | 1.3 ($\pm 0.7$) |
| ER chaperones | Calnexin (CNX) | 1.0 | 0.9 ($\pm 0.2$) | 1.2 ($\pm 0.6$) | 1.2 ($\pm 0.5$) |
| | Calreticulin (CRT) | 1.0 | 1.6 ($\pm 0.5$)# | 0.9 ($\pm 0.3$) | 0.8 ($\pm 0.3$) |
| NADPH generation | H6PD | 1.0 | 2.6 ($\pm 1.8$)* | 1.4 ($\pm 0.6$) | 1.4 ($\pm 0.6$) |

**Fig 1.**
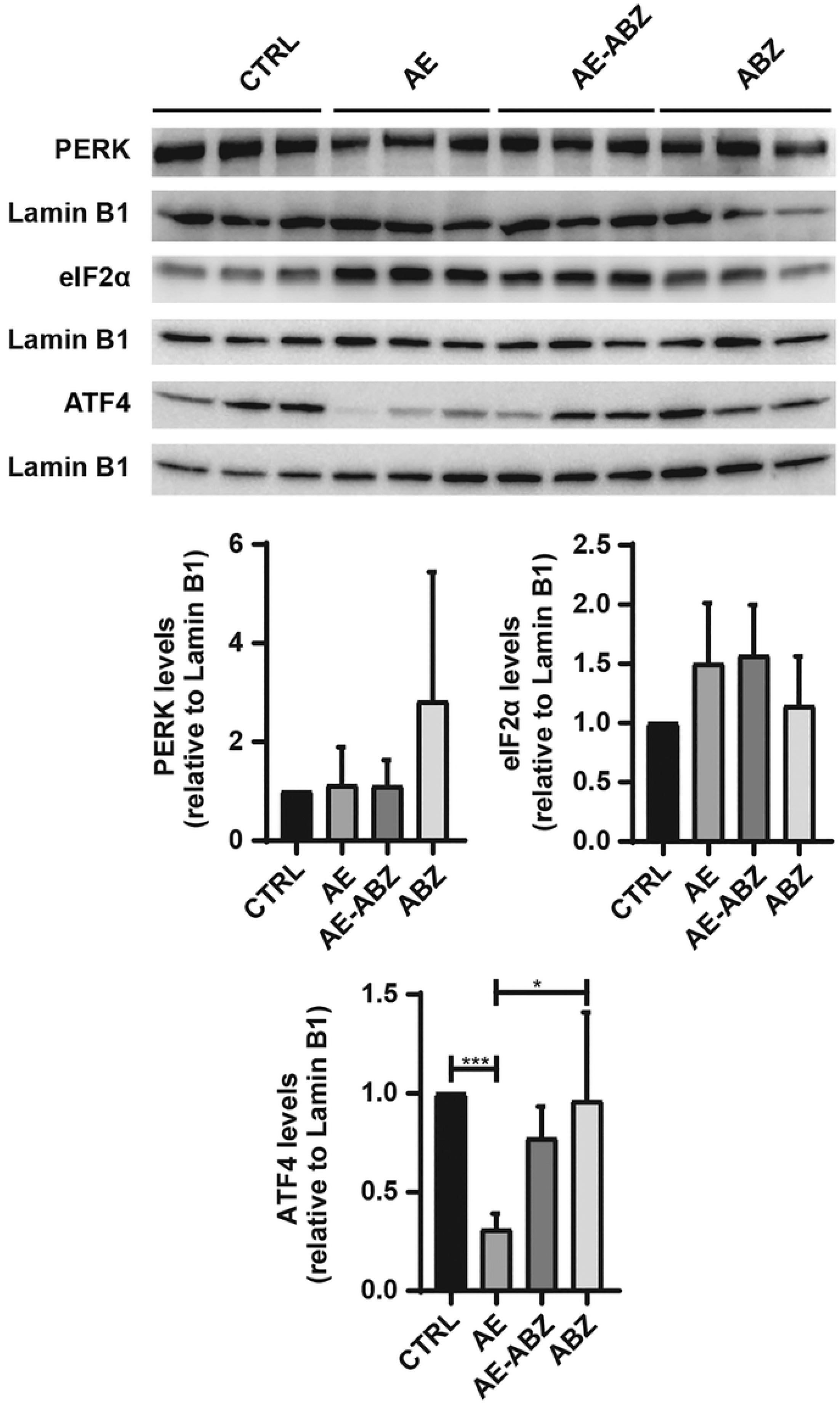
*E. multilocularis* infection decreases ATF4 protein levels. Western blot and semi-quantitative analysis by densitometry of protein levels of PERK, eIF2α and ATF4 in mock-infected control mice (CTRL), *E. multilocularis* infected mice (AE), infected mice treated with ABZ (AE-ABZ) or uninfected mice treated with ABZ (ABZ). One representative blot (of two) containing samples from three different mice is shown in the top panel. Densitometry results represent data from the two blots on samples from six mice (mean ± SD), normalized to lamin B1 control and with CTRL set as 1. No outliers were excluded. The non-parametric Kruskal-Wallis test was used to assess significance. *P≤0.05; ***p≤ 0.001.

**Fig 2.**
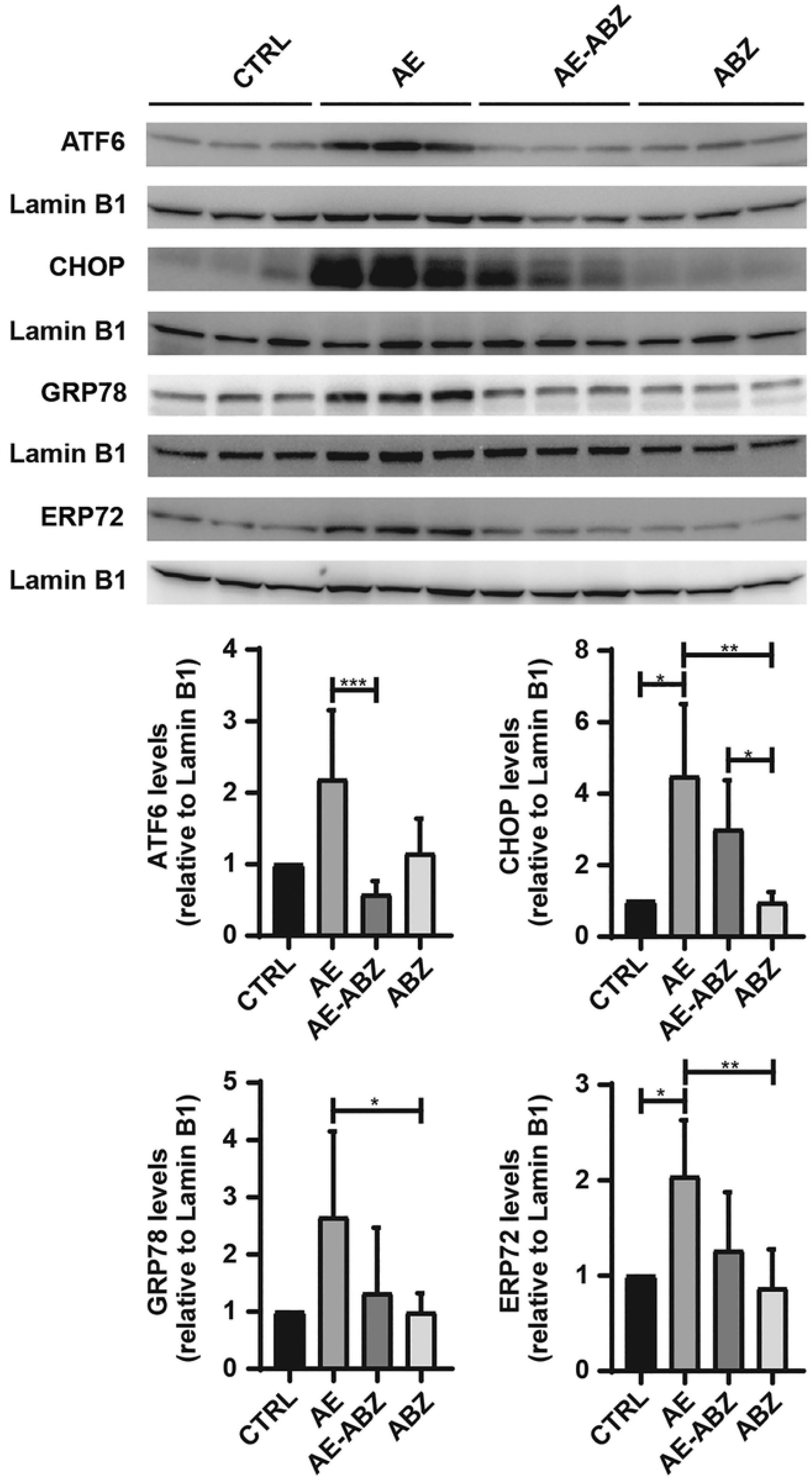
Induction of the ATF6-branch of the UPR by *E. multilocularis* infection. Western blot and semi-quantitative analysis by densitometry of the protein levels of ATF6, CHOP, GRP78 and ERP72 in mock-infected control mice (CTRL), *E. multilocularis* infected mice (AE), infected mice treated with ABZ (AE-ABZ) or uninfected mice treated with ABZ (ABZ). One representative blot (of two) containing samples from three different mice is shown in the top panel. Densitometry results represent data from the two blots on samples from six mice (mean ± SD), normalized to lamin B1 control and with CTRL set as 1. No outliers were excluded. The non-parametric Kruskal-Wallis test was used to assess significance. *P≤0.05; **p≤0.01; ***p≤ 0.001.

**Fig 3.**
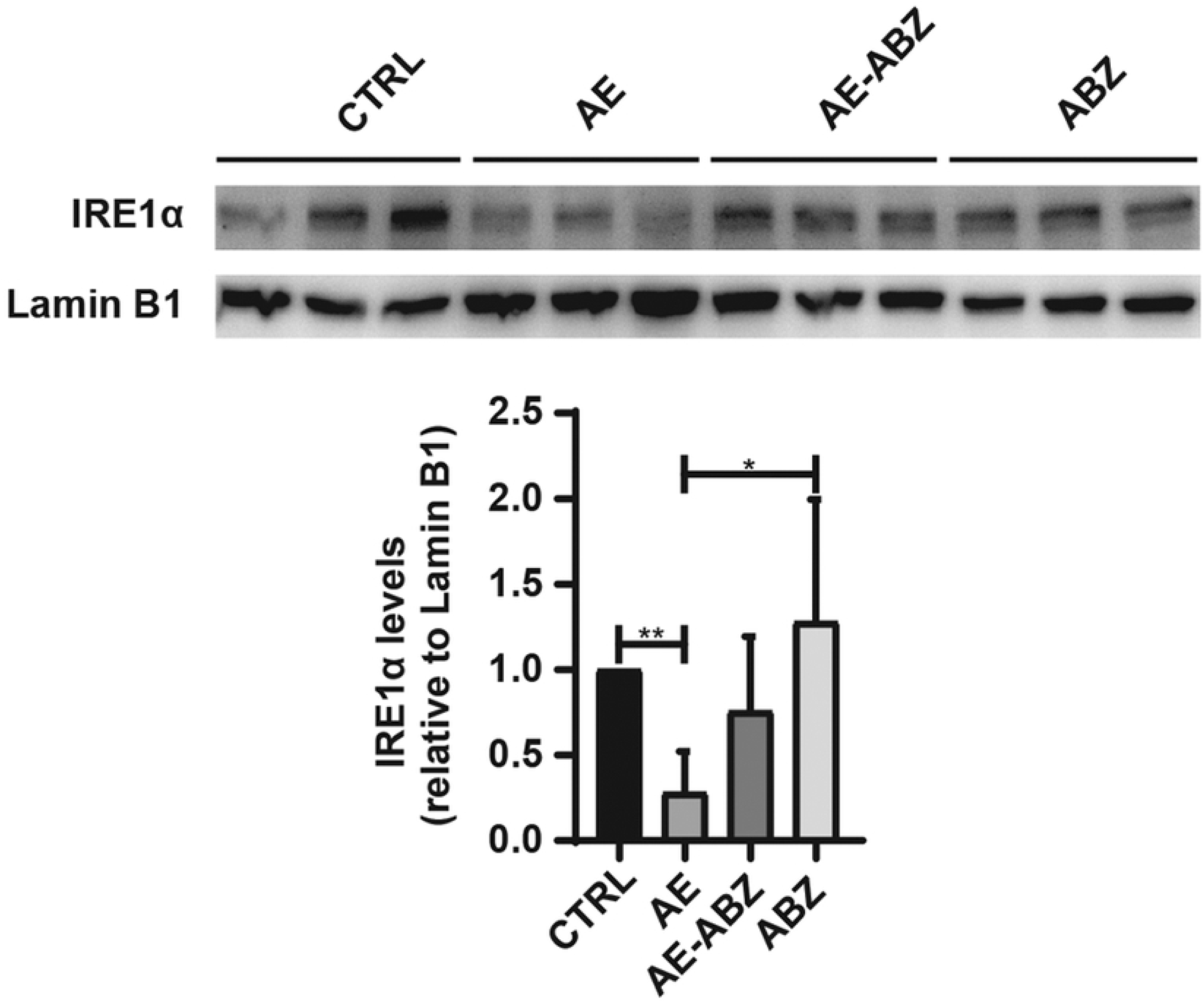
Decreased IRE1α protein expression upon *E. multilocularis* infection. Western blot and semi-quantitative analysis by densitometry of IRE1α protein levels in mock-infected control mice (CTRL), *E. multilocularis* infected mice (AE), infected mice treated with ABZ (AE-ABZ) or uninfected mice treated with ABZ (ABZ). One representative blot (of two) containing samples from three different mice is shown in the top panel. Densitometry results represent data from the two blots on samples from six mice (mean ± SD), normalized to lamin B1 control and with CTRL set as 1. No outliers were excluded. The non-parametric Kruskal-Wallis test was used to assess significance. *P≤0.05; **p≤0.01.

Additional proteins with a role in ER-redox regulation and ERS include the ER resident lectin chaperones CNX and CRT. Whilst CNX protein levels were unaffected by *E. multilocularis* infection, CRT protein expression was significantly increased in AE mice compared to controls (Table 1, Fig 4). Additionally, the expression levels of the luminal NADPH-generating enzyme H6PD were determined, revealing a 2.6-fold higher expression in AE compared to control mice (Table 1, Fig 5).

**Fig 4.**
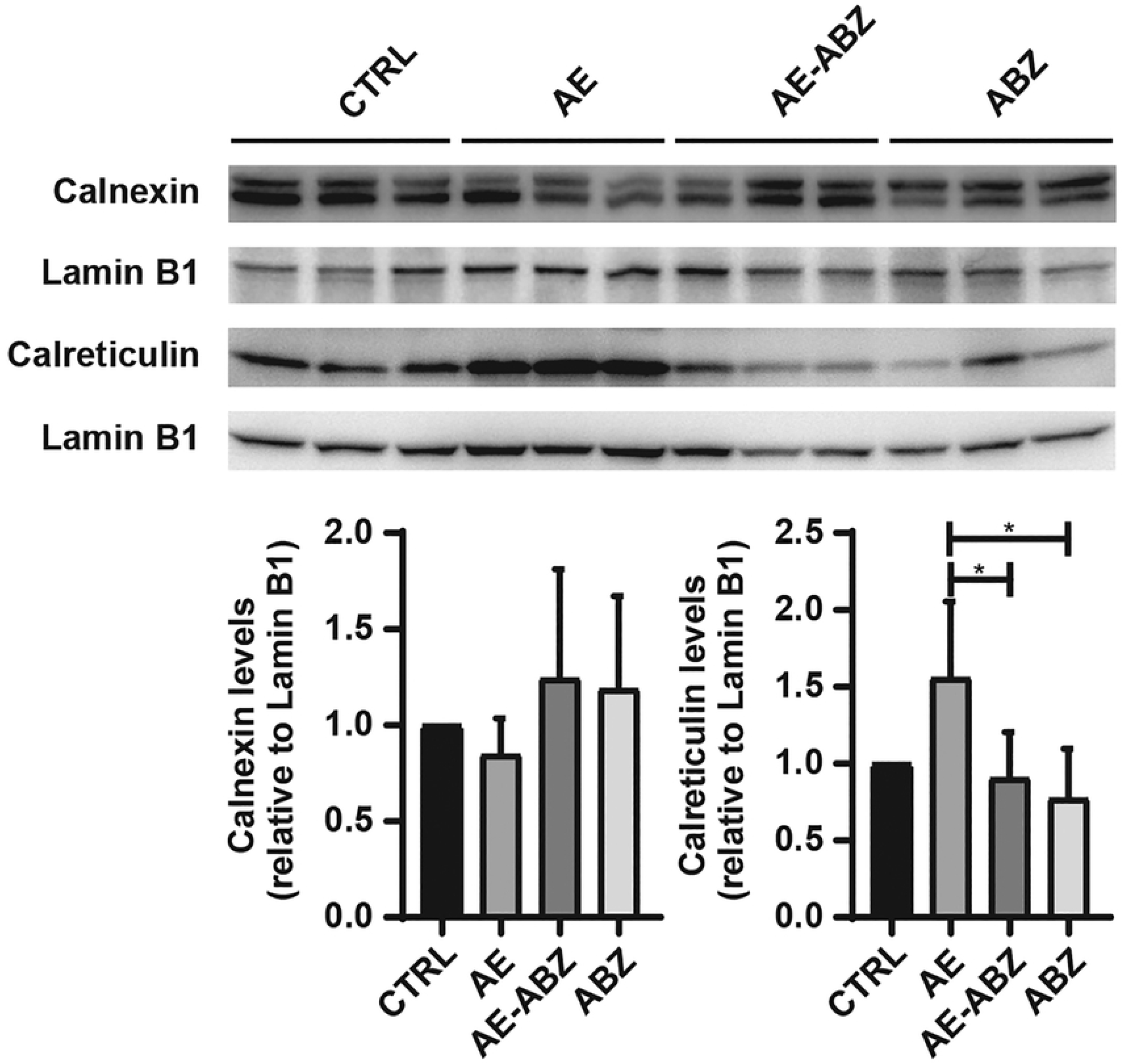
The expression of the luminal chaperone CRT is increased upon *E. multilocularis* infection and reversed by ABZ treatment. Western blot and semi-quantitative analysis by densitometry of CNX and CRT protein levels in mock-infected control mice (CTRL), *E. multilocularis* infected mice (AE), infected mice treated with ABZ (AE-ABZ) or uninfected mice treated with ABZ (ABZ). One representative blot (of two) containing samples from three different mice is shown in the top panel. Densitometry results represent data from the two blots on samples from six mice (mean ± SD), normalized to lamin B1 control and with CTRL set as 1. No outliers were excluded. The non-parametric Kruskal-Wallis test was used to assess significance. *P≤0.05.

**Fig 5.**
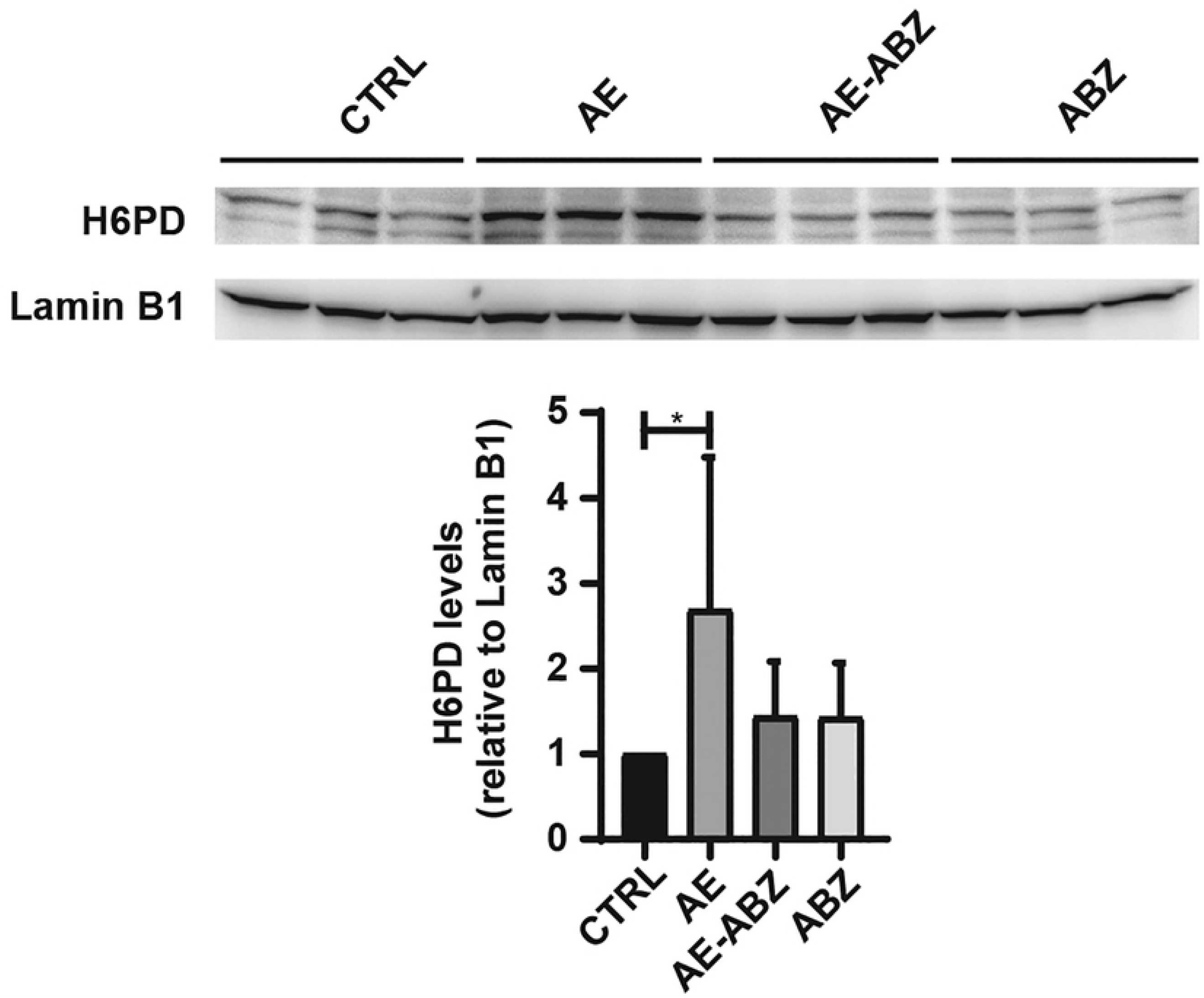
Increased H6PD protein expression upon *E. multilocularis* infection. Western blot and semi-quantitative analysis by densitometry of protein levels of H6PD in mock-infected control mice (CTRL), *E. multilocularis* infected mice (AE), infected mice treated with ABZ (AE-ABZ) or uninfected mice treated with ABZ (ABZ). One representative blot (of two) containing samples from three different mice is shown in the top panel. Densitometry results represent data from the two blots on samples from six mice (mean ± SD), normalized to lamin B1 control and with CTRL set as 1. No outliers were excluded. The non-parametric Kruskal-Wallis test was used to assess significance. *P≤0.05.

### Treatment with ABZ reverses the effects of AE on proteins involved in UPR and ERS

Treatment of AE mice with ABZ (AE-ABZ group) resulted in a reversal of the *E. multilocularis* induced alterations of UPR and ERS related protein expression (Table 1). Also the effects on the ER chaperones CNX and the NADPH-generating H6PD were reversed by ABZ treatment. An exception was CHOP that was still upregulated in ABZ treated infected mice. Importantly, ABZ did not cause any significant alterations in the expression of the proteins analyzed in uninfected control mice (Table 1, Figs 1-5). Protein levels of PERK show a trend to be increased in ABZ treated, uninfected animals (Fig 1); however, this did not reach significance due to high variance in the detected signals.

### Increased miR-146a-5p and miR-1839-5p expression in secondary *E. multilocularis* infection and reversal by ABZ treatment

Boubaker et al. [59], using an early stage mouse model of *E. multilocularis* infection, identified several miRs with altered expression in liver tissues from infected mice. In the present study, the levels of the miRs exhibiting a target site in the 3’UTR of genes involved in UPR and ERS pathways were determined. The analysis of the seven mouse miRs miR-148a-3p, miR-15a-5p, miR-22, miR-146a-5p, miR-1839-5p, miR-30a-5p and miR-30a-3p revealed significantly higher levels (2.1-fold and 3.2-fold, respectively) of miR-1839-5p and miR-146a-5p in liver tissue samples of *E. multilocularis* infected mice (AE) compared to control animals (CTRL). The other miRs remained unchanged (S3 Fig). Interestingly, ABZ treatment of AE mice decreased miR-1839-5p (2.8 fold) and miR-146a-5p (5.8 fold) compared to the levels found in CTRL animals or even lower, and ABZ alone tended to decrease miR-1839-5p and miR-146a-5p expression levels (Fig 6).

**Figure 6.**
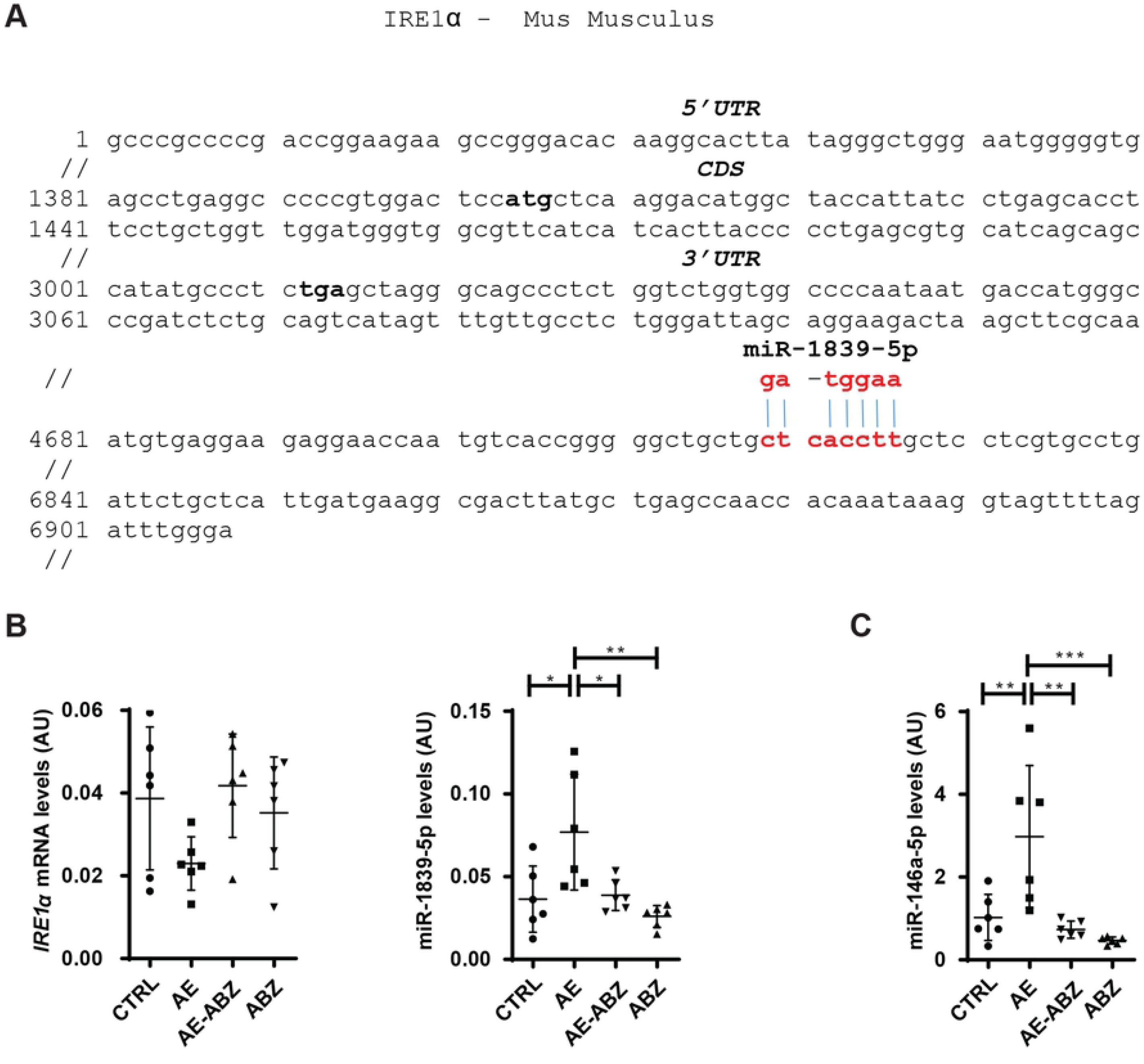
Altered expression of IRE1α related miRs upon *E. multilocularis* infection and reversal by ABZ treatment. A) Nucleotide sequence containing the murine IRE1α mRNA 3’-UTR. The start and stop codon of the IRE1α CDS are indicated in bold and the miR-1839-5p binding site is highlighted by red and bold letters. B) IRE1α mRNA levels normalized to the β-actin housekeeping gene and relative to the levels obtained in control mice, and miR-1839-5p levels normalized to the Sno234 housekeeping gene and relative to uninfected controls. C) miR-146-5p levels normalized to Sno234 and relative to uninfected controls. B, C) mock-infected control mice (CTRL _n=6_), *E. multilocularis* infected mice (AE _n=6_), infected mice treated with ABZ (AE-ABZ _n=6_) or uninfected mice treated with ABZ (ABZ _n=6_). Results represent mean ± SD. No outliers were excluded. One-way ANOVA test was used to assess significance. *P≤0.05; **p≤0.01; ***p≤ 0.001.

## Discussion

Recent studies on viral, bacterial or intracellular parasitic infections emphasize the important roles of the UPR and ERS pathways in pathogen induced diseases [23-25, 64, 65]. Activation of the UPR, a specific form of ERS triggered by an accumulation of unfolded or misfolded proteins within the ER, can be mediated by three branches, represented by the ER transmembrane stress sensor proteins ATF6, PERK and IRE1α [66-74] (Fig 7). In non-stressed cells, these proteins remain in an inactive state, bound to the luminal chaperone GRP78. Upon activation, GRP78 is released to support luminal protein folding, followed by the activation of ATF6, PERK and IRE1α and their downstream targets such as eIF2α, ATF4, XBP1 and CHOP in order to mediate the stress response [75-77].

**Fig 7.**
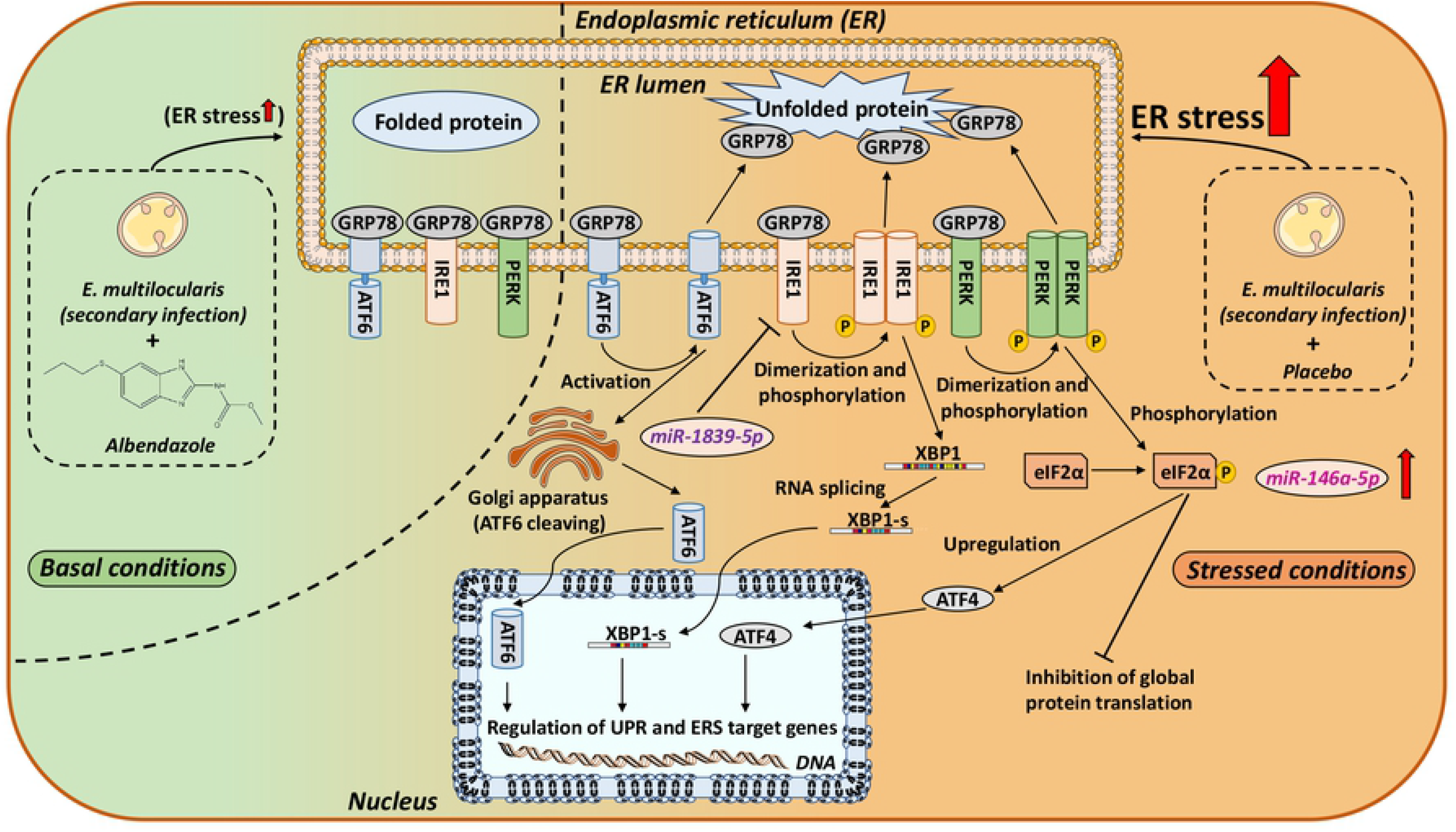
Schematic overview of ERS signaling pathways under basal and *E. multilocularis* infection stressed conditions. The ER chaperone GRP78 binds to unfolded luminal proteins and dissociates from the three major ERS sensors: ATF6, IRE1α and PERK. Loss of GRP78 binding leads to the translocation of ATF6 to the golgi apparatus, where it is cleaved by proteases. The cleaved form of ATF6 translocates into the nucleus to act as a transcription factor for ER chaperons (*e*.*g*. ERP72) and ERS related genes. ERS promotes IRE1α dimerization and autophosphorylation, which activates the endoribonuclease activity resulting in the splicing and thereby activation of XBP. XBP1-s promotes the expression of ERAD related genes and chaperones (*e*.*g*. GRP78). Activation of PERK is initiated by dimerization and self-phosphorylation. Activated PERK phosphorylates eIF2α, leading to eIF2α-mediated inhibition of global protein translation in order to decrease the luminal protein load. Besides, phosphorylated eIF2α increases the transcription of ATF4, which in turn upregulates expression of genes related to cell homeostasis restoration. If prolonged ERS occurs and pro-adaptive UPR fails, ATF4 induces genes (including CHOP) leading to apoptosis. During ERS, increased levels of miR-1839-5p are proposed to control IRE1α gene expression and therefore affect the cellular ERS response.

The results of the present study revealed a pronounced induction of ATF6 in livers of mice infected with *E. multilocularis*. In contrast, the PERK and IRE1α branches were not activated but rather down regulated. Unfortunately, the levels of phosphorylated PERK, eIF2α and IRE1α could not be assessed due to the failure to identify specific antibodies. The decreased levels of ATF4 in livers of infected animals suggests that the observed upregulation of CHOP is caused by enhanced ATF6 activity. CHOP, well-known as a mediator of apoptosis, was previously found to play an important role in the efficient expansion of the intracellular fungus *Histoplasma capsulatum* [78]. Following infection an increase in CHOP levels led to augmented apoptosis of macrophages, thus suppressing the host’s defense and contributing to the virulence of this particular pathogen. Another study, using intestinal epithelial cell lines, showed a direct effect of heat-labile enterotoxins of *Escherichia coli* on the induction upregulation of CHOP, which led to an accelerated apoptosis of the host cells [79]. Thus, the upregulation of CHOP in murine hepatocytes during *E. multilocularis* infection might similarly promote parasitic growth.

In contrast to the pro-apoptotic UPR mediator CHOP, the protein levels of the PERK target ATF4 were significantly decreased in livers of *E. multilocularis* infected compared to mock-infected mice. This is different from a previous study on human cutaneous leishmaniasis where both CHOP and ATF4 were found to be upregulated [80]. Decreased levels of ATF4 were recently described as a mechanism of acquired resistance to cope with a limited availability of amino acids in cancer cells [81]. Unrestricted tumor growth requires a high demand of nutrients and has been associated with a depletion of essential amino acids in the tumor tissue. Similar metabolic perturbations and adaptive responses may occur in patients with hepatic AE. A recent study summarizing analyses of serum samples from *E. multilocularis* infected and healthy adults (group size: n=18) revealed decreased levels of branched-chain amino acids such as leucine, isoleucine and valine along with lowered levels of serine and glutamine in samples from infected patients [82]. In contrast, the aromatic amino acids tyrosine and phenylalanine were increased, together with glutamate. Thus, the observed decrease in ATF4 expression may be a response to adapt the amino acid availability in the situation of parasitic growth.

Similar to ARF4 also IRE1α protein expression levels were decreased in liver tissues of AE infected mice. The reason of the decreased IRE1α expression in *E. multilocularis* infected mice and the underlying mechanism remain unclear. IRE1 enzymes are transmembrane proteins exhibiting Ser/Thr protein kinase and endoribonuclease activities and acting as major ERS sensors [83, 84]. There are two IRE1 isoforms in mammals: the ubiquitously expressed IRE1α and IRE1β which is predominantly expressed in the intestine and lung [85]. Further analysis of the liver resident IRE1α showed that the decreased protein expression in *E. multilocularis* infected mouse livers is supported by lower mRNA levels along with an increased expression of miR-1839-5p that has a target site in the 3’UTR of IRE1α as predicted by the computer-based programs Targetscan (Whitehead Institute, Cambridge, MA, USA, RRID:SCR_010845) [86] and RNA22 (Thomas Jefferson University, Philadelphia, PA, USA, RRID:SCR_016507) [87]. Additionally, miR-146a-5p was found to be enhanced in livers of infected mice. An earlier study in primary dermal fibroblasts provided evidence for a down regulation of miR-146a-5p by IRE1-dependent cleavage in response to UPR activation. A decreased hepatic IRE1α expression and activity upon *E. multilocularis* infection might suppress the activation of pro-inflammatory cytokines such as IL-1β as well as of NF-κB. Whether this promotes the progression of AE remains to be investigated.

An extensive analysis of miRs altered in livers of mice after primary infection with *E. multilocularis* by Boubaker *et al*. identified, besides miR-1839-5p and miR-146a-5p, several other miRs that were dysregulated, *i*.*e*. miR-148-3p, miR-15a-5p, miR-22, miR-30a-5p and miR-30a-3p [59]. The fact that miR-1839-5p and miR-146a-5p were increased in primary as well as in the secondary form of infection suggests these two miRs as potential biomarkers of AE. In this regard, Luis et al. reported an association of several circulating miRs, including miR-146a-5p, with ERS and organ damage in a model of trauma hemorrhagic shock [88].

Further, Wilczynski et al. reported increased miR-146a expression levels in tumor tissues of patients with ovarian cancer [89]. The advanced AE resembles a tumorigenic situation with alterations in the microenvironment and immune responses. Thus, follow-on research should address whether miR-146a-5p and miR-1839-5p can serve as serum biomarkers of AE.

Beside the UPR, the ER-associated degradation (ERAD) is an important quality control machinery to cope with ER stressors. ERAD plays a crucial role in the degradation of terminally misfolded proteins by retro-translocating them from the ER to the cytoplasm for deglycosylation and ubiquitination and subsequent proteasomal degradation [90, 91]. Prior to ERAD, misfolded proteins undergo repeated cycles of re-folding by the assistance of several ER-resident chaperones including lectins such as CRT and CNX, protein disulfide isomerase family members like ERP72 and ERP57 as well as members of the heat shock protein 70 family (e.g. GRP78) [92-95]. The elevated expression of CRT together with GRP78 and ERP72 indicates a higher demand for protein folding capacity in the ER in livers from infected mice. This was accompanied by an elevated demand for NADPH redox equivalents in the ER and/or an enhanced need for the products of the ER pentose phosphate pathway as indicated by the elevated H6PD expression. H6PD was found to promote cancer cell proliferation and the modulation of its expression affected GRP78, ATF6 and CHOP, emphasizing its role in ERS regulation [96].

Importantly, treatment with the parasitostatic benzimidazole ABZ, which was shown to decrease the weight of parasitic cysts in the livers of *E. multilocularis* infected mice, reversed the observed effects on UPR and ERS pathways and on associated ERAD and ER redox genes. In the absence of infection, ABZ did not affect any of the investigated ER related targets, underlining its favorable safety profile regarding ERS related adverse effects.

In conclusion, the present study showed that *E. multilocularis* infection leads to a modulation of the UPR, characterized by an activation of the ATF6-branch with an upregulation of CHOP along with decreased ARF4 and IRE1α protein levels and increased miR-1839-5p and miR-146a-5p that could serve as potential biomarkers of *E. multilocularis* infections. ABZ, the most commonly used drug to treat AE ameliorated the effects of *E. multilocularis* infection on ER related genes. Whether drugs targeting UPR and ERS pathways in combination with ABZ may improve the treatment of AE remains to be explored.

## Supplementary data

**S1 Table. Antibodies and corresponding dilutions**.

**S2 Table. Primers used for RT-qPCR**.

**S1 Fig. Schematic overview of experimental setup**. Animals were divided into four groups: CTRL_(n=6)_, AE_(n=6)_, AE-ABZ_(n=6)_, and ABZ_(n=6)_. CTRL and ABZ mice received an intraperitoneal administration of 100 µL PBS. AE and AE-ABZ mice were treated by secondary *E. multilocularis* infection using approximately 100 vesicular cysts resuspended in 100 µL PBS. Treatment started 6 weeks after infection. CTRL and AE mice received 100 µL corn oil orally 5 times per week for 8 weeks. AE-ABZ and ABZ mice received ABZ (200 mg/kg body weight) in 100 µL corn oil orally 5 times per week for 8 weeks. Animals were sacrificed at the end of treatment.

**S2 Fig. XBP1 mRNA and XBP1-s mRNA levels are not altered upon *E. multilocularis* infection**. XBP1 and XBP1-s mRNA levels in mock-infected control mice (CTRL _n=6_), *E. multilocularis* infected mice (AE _n=6_), infected mice treated with ABZ (AE-ABZ _n=6_) or uninfected mice treated with ABZ (ABZ _n=6_). Results represent mean ± SD. No outliers were excluded. One-way ANOVA was applied to test significance.

**S3 Fig. *E. multilocularis* infection does not affect miR-15a-5p, miR-148a-3p, miR-22-3p, miR-30a-3p and miR-30a-5p expression levels**. miR-15a-5p, miR-148a-3p, miR-22-3p, miR-30a-5p and miR-30a-3p levels, in mock-infected, mock-treated mice (CTRL _n=6_) and *E. multilocularis* infected mock-treated mice (AE _n=6_). Results represent mean ± SD. No outliers were excluded. Two-tailed unpaired t-test was applied to test significance.

**S1 File. Raw data of western blots used to produce graphs and figures**.

**Striking image**. Experimental setup and key finding that *E. multilocularis* infection causes ER-stress in mouse liver. Treatment of infected mice using the anthelminthic drug albendazole attenuates the altered expression of selected ER-stress related genes.

## Acknowledgements

None.

## Data Availability

All relevant data are within the manuscript and in its supporting information files.

## Author Contributions

***Conceptualization*** Alex Odermatt, Michael Weingartner, Fadi Jebbawi.

***Data curation*** Michael Weingartner, Fadi Jebbawi, Simon Stücheli

***Formal analysi*s** Michael Weingartner, Fadi Jebbawi

***Funding acquisition*** Alex Odermatt, Britta Lundström-Stadelmann

***Investigation*** Michael Weingartner, Fadi Jebbawi, Junhua Wang

***Methodology*** Michael Weingartner, Fadi Jebbawi, Junhua Wang, Britta Lundström-Stadelmann

***Project administration*** Michael Weingartner, Fadi Jebbawi

***Resources*** Alex Odermatt, Britta Lundström-Stadelmann, Bruno Gottstein, Guido Beldi

***Supervision*** Alex Odermatt, Britta Lundström-Stadelmann, Bruno Gottstein, Guido Beldi

***Visualization*:** Michael Weingartner

***Writing – original draft*:** Michael Weingartner

***Writing – review & editing*** All authors

## Rights and permissions

Presented work is licensed under a Creative Commons Attribution 3.0 (CC BY 3.0) International License. The images or other third-party material in this article are included in the article’s Creative Commons license. Unless indicated otherwise; if the material is not included under the Creative Commons license, users will need to obtain permission from the license holder to reproduce the material. To view a copy of this license, visit: https://creativecommons.org/licenses/by/3.0/

## Abbreviations

ABZ: albendazole
AE: alveolar echinococcosis
ATF4: activating transcription factor 4
ATF6: activating transcription factor 6
CHOP: CCAAT/enhancer-binding protein homologous protein
CNX: calnexin
CRT: calreticulin
CTRL: control
*E. multilocularis*: *Echinococcus multilocularis*
GRP78: glucose-binding protein 78
eIF2α: eukaryotic initiation factor 2α
ER: endoplasmic reticulum
ERAD: ER-associated degradation
ERP72: endoplasmic reticulum resident protein 72
ERS: endoplasmic reticulum stress
H6PD: hexose-6-phosphate dehydrogenase
HRP: horse-radish peroxidase
IRE1α: inositol-requiring enzyme 1α
microRNA(s): miR(s)
PERK: protein kinase R (PKR) like ER kinase
PVDF: polyvinylidene difluoride
RIPA: radioimmunoprecipitation assay
RT-qPCR: real-time quantitative polymerase chain reaction
UPR: unfolded protein response
XBP1(-s): X-box binding protein 1 (-spliced)

